# Integration and multiplexing of positional and contextual information by the hippocampal network

**DOI:** 10.1101/269340

**Authors:** Lorenzo Posani, Simona Cocco, Rémi Monasson

## Abstract

The hippocampus is known to store cognitive representations, or maps, that encode both positional and contextual information, critical for episodic memories and functional behavior. How path integration and contextual cues are dynamically combined and processed by the hippocampus to maintain these representations accurate over time remains unclear. To answer this question, we propose a two-way data analysis and modeling approach to CA3 multi-electrode recordings of a moving rat submitted to rapid changes of contextual (light) cues, triggering back-and-forth instabitilies between two cognitive representations (Jezek et al, Nature 478, p 246 (2011)). We develop a dual neural activity decoder, capable of independently identifying the recalled cognitive map at high temporal resolution (comparable to theta cycle) and the position of the rodent given a map. Remarkably, position can be reconstructed at any time with an accuracy comparable to fixed-context periods, even during highly unstable periods. These findings provide evidence for the capability of the hippocampal neural activity to maintain an accurate encoding of spatial and contextual variables, while one of these variables undergoes rapid changes independently of the other. To explain this result we introduce an attractor neural network model for the hippocampal activity that process inputs from external cues and the path integrator. Our model allows us to make predictions on the frequency of the cognitive map instability, its duration, and the detailed nature of the place-cell population activity, which are validated by a further analysis of the data. Our work therefore sheds light on the mechanisms by which the hippocampal network achieves and updates multi-dimensional neural representations from various input streams.

**Author summary:** As an animal moves in space and receives external sensory inputs, it must dynamically maintain the representations of its position and environment at all times. How the hippocampus, the brain area crucial for spatial representations, achieves this task, and manages possible conflicts between different inputs remains unclear. We propose here a comprehensive attractor neural network-based model of the hippocampus and of its multiple input streams (including self-motion). We show that this model is capable of maintaining faithful representations of positional and contextual information, and resolves conflicts by adapting internal representations to match external cues. Model predictions are confirmed by the detailed analysis of hippocampal recordings of a rat submitted to quickly varying and conflicting contextual inputs.

## 1. Introduction

Following the discovery of place cells, which specifically fire at determined positions in space [1], the hippocampus was recognized as an essential brain area for spatial representations and memories. These cognitive representations, or maps, actually code for more than position in physical space, and are also strongly informative about context [2], including physical features of the background, such as visual landmarks, light, odors, auditory stimuli, as well as more abstract conditions, such as the emotional state or the task to be performed [3–8].

A fundamental property of the hippocampus is its capacity to memorize multiple cognitive maps [1, 9–11]. This property may result from specific recurrent synaptic connectivity in the hippocampal CA3 region [12, 13], and can be theoretically understood in the framework of continuous attractor neural networks (CANN) [14, 15]. Thanks to the remapping properties of place cells, multiple maps can be memorized in the same connectivity matrix with almost no interference between them [16–19].

Cognitive maps may be retrieved when the animal explores again the corresponding environments, or be quickly and intermittently recalled depending on the most relevant behavorial information at that moment [20]. Different sources of inputs to the hippocampus concur to form, recall, and dynamically maintain cognitive maps [21]. Changes in visual cues and landmarks may substantially affect place field shape and positioning [5]. The Path Integrator (PI), capable of integrating proprioceptive, vestibular and visual flow inputs and possibly supported by the grid-cell network in the medial-enthorinal cortex (mEC) [22], allows the animal to update the neural representation during navigation [23]. The path integrator is itself sensitive to other sources of inputs, and undergoes reset in case of large disagreement with external landmarks or sensory information [24].

Insights about how these different inputs contribute to hippocampal representations were recently obtained by studying the effects of mismatches between path-integration and visual sensory information, in particular in virtual reality settings [25, 26]. In another study Jezek et al showed how abrupt changes in visual context (light conditions) during active exploration by a rodent resulted in fast flickering between context-associated maps in CA3 on the theta time scale [27] (Fig. 1A). Though they are largely artificial, these conditions offer a rare window on the fast dynamics of the place-cell population, and on how this dynamics is shaped by the various inputs.

**Fig. 1:**
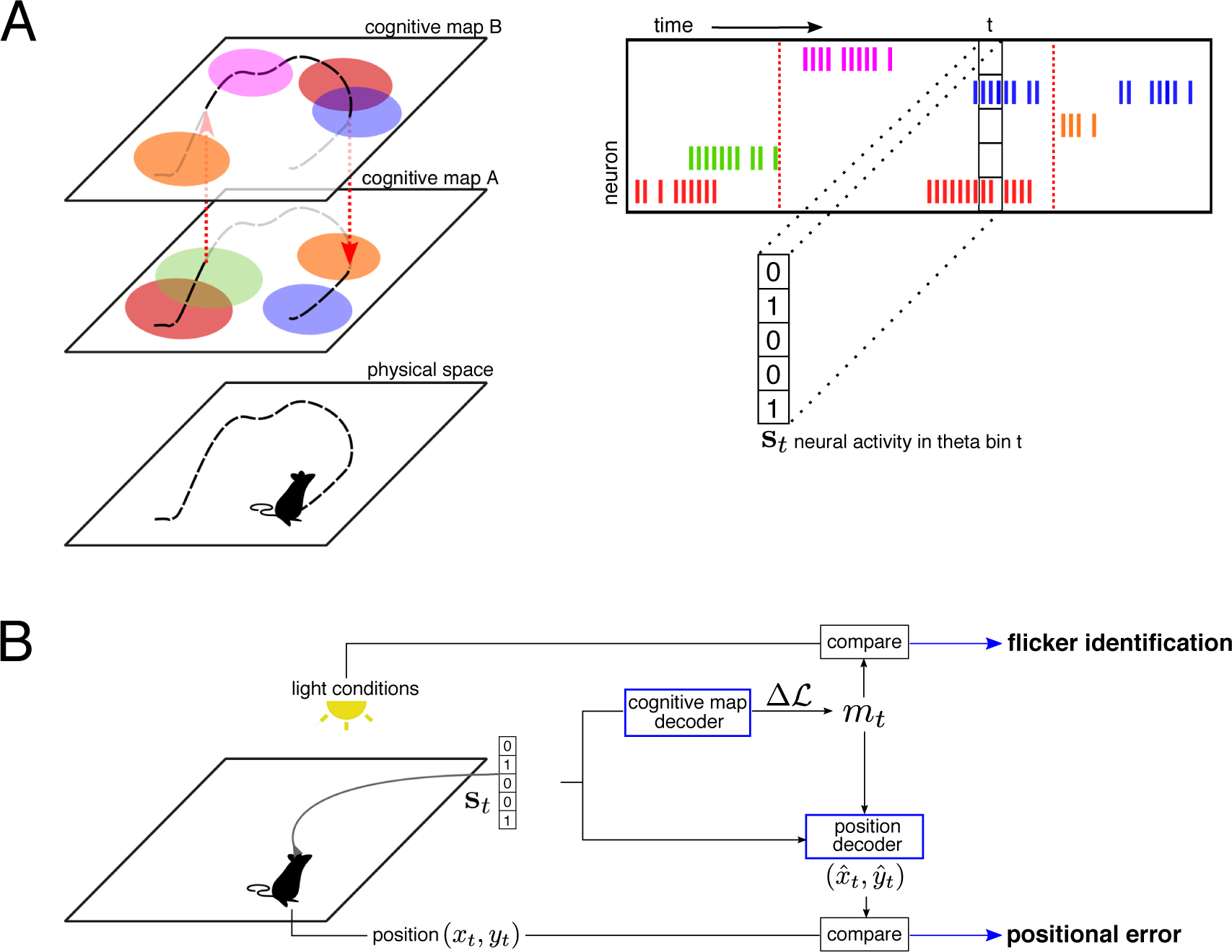
Neural encoding and decoding of cognitive maps and position. **A. Schematic description of Jezek et al’s experiment.** As a rodent is moving in an environment, its position **r** = (*x, y*) is tracked over time, and a population of place cells is recorded (see raster plot). The activity of each cell is then binarized (0: silent cell, 1: active cell) to define the activity pattern s_*t*_ in theta cycle *t*. One out of two cognitive maps, established during the training sessions, is recalled at any time; change of maps are located by vertical dashed arrows. Place cells may have place fields in both cognitive maps (red, orange, and blue cells) or in one map only (green and purple cells). In addition, pair of neurons may be active simultaneously or not depending on the map; for instance the red and blue neurons have overlapping place fields in map *B*, but not in *A*. Hence, pairwise correlations are a fingerprint of the map. **B. Sketch of the dual decoder.** The neural activity alone is used to decode the retrieved map *m*_*t*_ as a function of time, and then to infer the position of the animal based on the place fields in the decoded cognitive map. Mismatches between the decoded maps and the external light cues define flickers over time. The distance between the predicted and real positions defines the positional error *ϵ*_*t*_.

Despite these studies, how contextual and PI inputs are combined by the hippocampal network to produce cognitive maps and accurate positional encoding is not fully understood yet. In this work, we carefully reanalyze and model the experiment of Jezek et al to address this issue. We first introduce of a dual inference method capable of extracting reliably and independently the encoded map [28] and the encoded position [29, 30] from the recorded spiking activity alone (Fig. 1B). Our dual decoder allows us to robustly show that the hippocampal activity always encodes the correct location in the retrieved map, even during the fast, unstable dynamics of the cognitive maps, as put forward in [27].

To explain this robust encoding, we propose a CANN model of the hippocampal circuitry, capable of storing multiple cognitive maps; the model is fed by visual-cue and path-integration inputs projecting on the place-cell populations supporting those maps [31]. The path integrator is, in turn, influenced by the hippocampal activity, closing an interaction loop between the hippocampus and the mEC [32]. Our model not only reproduces the flickering phenomenology and the stable encoding of position, but also makes several precise predictions on the dynamics of cognitive maps, the relative strength of inputs, and the intricate activation of place-cell populations supporting the two maps. These predictions are corroborated by a further detailed analysis of Jezek et al’s data. Our work therefore proposes explicit mechanisms by which the hippocampus could be capable of encoding various contextual and self-locomotion information in multi-dimensional representations, and of updating them accurately on fast time scales.

## 2 Results

Jezek et al trained a rodent in two environments (square boxes), equal in size and shape, but differing by their light conditions [27]. A population of 34 CA3 place cells was recorded during reference sessions with fixed light conditions, and shown to define environment-specific maps, denoted by *A* and *B*. In a subsequent test session, taking place in a single box, instantaneous switches between environmental light conditions triggered the instability of the recalled cognitive map, which flickered back and forth between the two corresponding environments.

The neural activity s_*t*_ of the population in any theta cycle *t* encodes information on the context (the set of rules that connects position to activity, i.e. the place fields defining the cognitive map) as well as on the specific position within the environment (Fig. 1A). We first introduce a dual decoder, able to independently infer the cognitive map and the position, at high temporal resolution. By comparing the inferred position to the true animal location, we then assess how precisely the position is represented in the population activity, irrespectively of the cognitive map in which it is neurally encoded (Fig. 1B).

## Functional network-based decoding of the cognitive map dynamics

Due to the global remapping properties of CA3, the intensities and mutual superpositions of place fields are specific to each environment (Fig. 1A). Consequently, the average firing rates and pairwise correlations of the place-cell population define a fingerprint of the corresponding cognitive map [33, 34]. We use the reference session recordings in each environment *m*(*A* or *B*) to compute this fingerprint statistics. We then build a model *P*_*m*_(s) that approximates the probability of observing the neural activity s when the cognitive map *m* is recalled. This model relies on the inference of a functional network of couplings between the place cells, reproducing the fingerprint statistics of map *m* [34, 35] (Methods). Given the activity s_*t*_ recorded in theta bin *t* during the test session, we then compare the two probabilistic models *P*_*a*_ and *P*_*B*_ to estimate which map m is more likely to have generated s_*t*_. The log-ratio

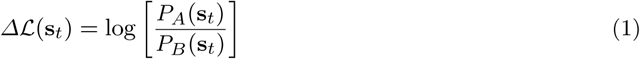

indicates whether the neural activity s_*t*_ is more similar to the neural patterns encountered in map *A* than to the ones of map *B* (large and positive *ΔL*), or typical of *B* and not of *A* (large in absolute value and negative *ΔL*). Comparing *ΔL* to a statistical significance threshold allows us to infer the map *m*_*t*_ (Methods). If the decoded map *m*_*t*_ is discordant with the imposed light conditions the theta bin is identified as a *flicker.*

As a control, we check that *ΔL* is mostly positive in reference sessions for environment *A* and negative for *B*, see Fig. 2A. Applying the decoder to the test session, we observe the presence of flickers, see Fig. 2B (yellow bins); flickers were first found in [27] with correlation-based methods requiring knowledge of the true position of the animal. An analysis of the temporal correlation of these flickers reveals that they typically persist over ~ 6 theta bins (Methods and Fig. 2C); hence, cognitive maps show some inertia extending beyond the theta scale.

**Fig. 2:**
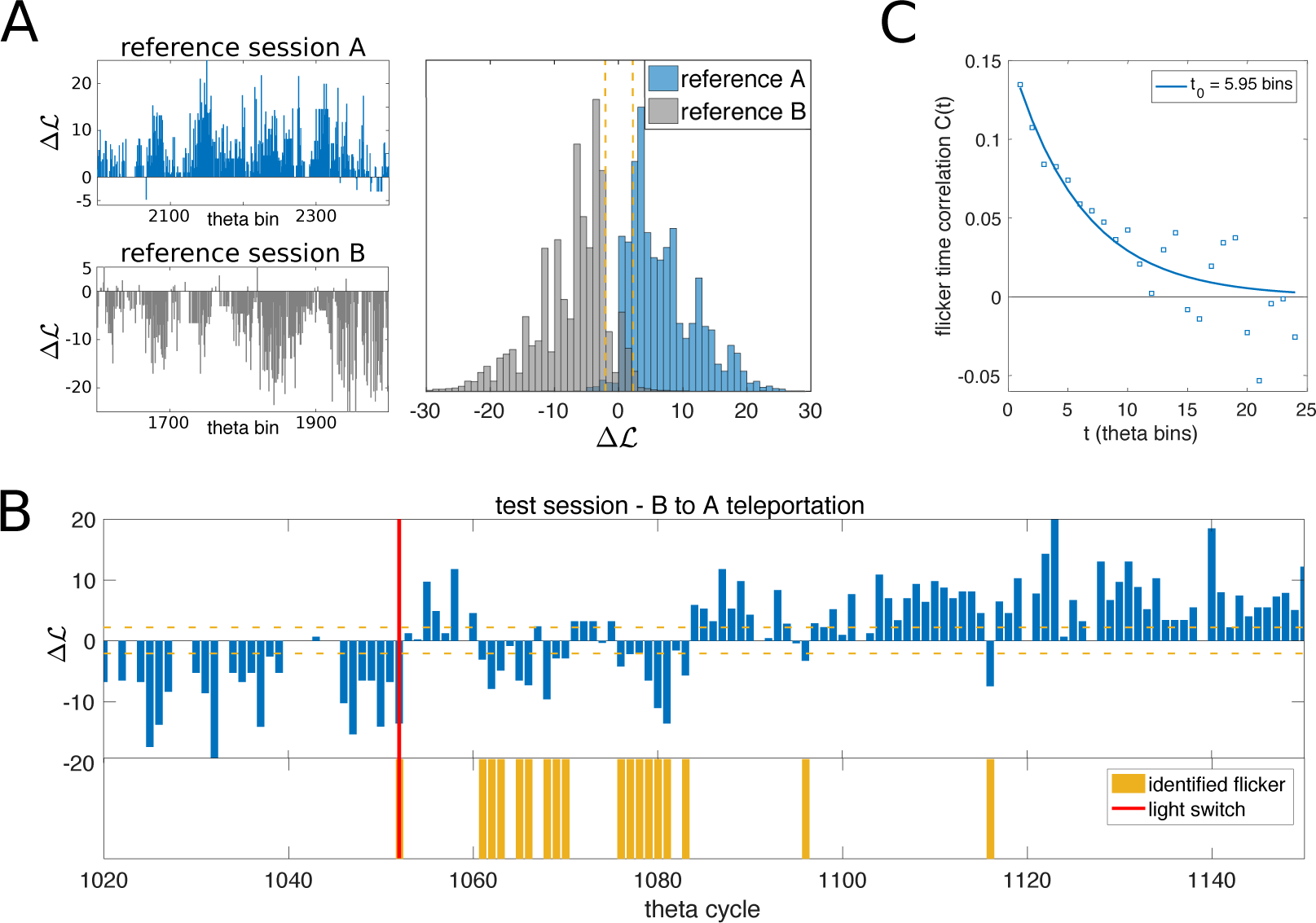
Decoding the cognitive map from neural activity. **A.** Map-decoding procedure applied to constant-environment reference sessions. The log-ratio *ΔL*_*t*_, Eqn. (1), which is the difference of likelihoods of map *A* and *B* given the recorded neural pattern in theta bin *t* (Methods), is mostly positive in the reference session with constant light-cue evoking *A* (blue) and mostly negative in the reference session *B* (grey). **B.** Time course of *ΔL* in a portion of the test session around one light switch (red vertical line). The emergence of flickers (disagreement between the recalled map and the light cue) is clearly visible after the light switch. Yellow horizontal dashed lines show the statistical threshold applied in map decoding and flicker identification (|*ΔL*| > *L*_0_ = log 10, Methods). Yellow bars represent the identified flicker instabilities, i.e. significative discordances *(L*_0_ *< − Lo*) between the decoded cognitive map (here, *B*) and the post-switch light conditions (here, *A*). **C.** time correlation of flickers, computed with significance threshold *L*_0_ = log 10. The exponential fit shows that correlations extend over ~ 6 theta bins, highlighting the tendency of the cognitive map to persist beyond the theta cycle.

## Position is accurately encoded even during flickering instabilities of the cognitive map

To assess if the fast dynamics of cognitive maps affects the quality of positional encoding we next reuse the neural activity pattern s_*t*_ in theta bin *t*, this time to infer the position of the animal. A naive Bayesian decoder [29, 30] takes as an input the above-decoded map *m*_*t*_ (Fig. 1B) and uses its place fields to estimate the position. The distance between the inferred position, 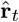, and the true position, **r**_*t*_, defines the positional error *ϵ*_*t*_. As shown in Fig. 3A, the positional error *ϵ*_*t*_ (blue line) is independent of the time elapsed after the light switch, and has a value comparable to the one obtained in fixed-environment conditions (blue dashed line). This result crucially depends on the fact that position is estimated according to the decoded map *m*_*t*_, which varies with time *t*. For comparison, in Fig. 3B we show the error if we decode the position according to the new, post-switch map (green line) or to the old, pre-switch one (red line) at all times. Both procedures result in similar, higher errors right after the light-switch, where flickers are frequent. The error with the post-switch map eventually decrease to fixed-environment value after few seconds, due to the rarity of flickers long after the light switch.

**Fig. 3:**
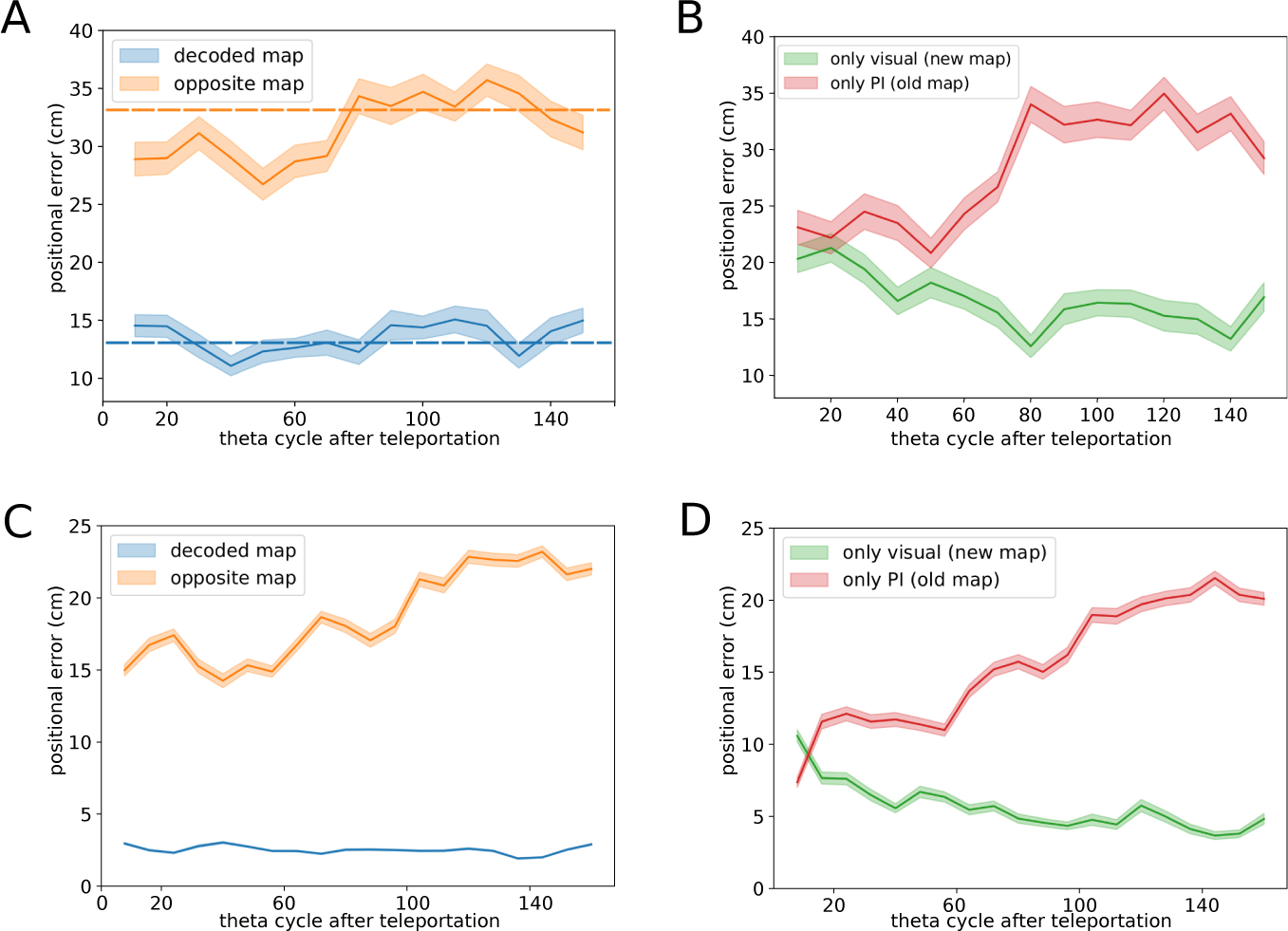
Positional error as a function of time from the teleportation light switch. **A. Recorded data. Positional error computed as a function of time from the teleportation (in units of theta cycle).** In each theta bin *t*, the map decoder is used to select which map *m*_*t*_ (set of place fields) to use to infer the position from the neural activity (Fig. 1A). When using the decoded map *m*_*t*_, the positional error (blue line) is comparable to the one found in constant-environment conditions (blue dashed line). The orange line shows the positional error if the opposite map (alternative to *m*_*t*_) is used for inferring position. The positional error is significantly reduced with respect to constant-environment conditions (orange dashed line) in the vicinity of the teleportation, and reaches comparable values after ~ 80 theta bins. Shaded areas represent the standard error computed over 15 teleportation events. **B. Recorded data**. Same as in panel A but using a fixed map of reference for position inference, irrespectively from the decoded map. Red and green lines show results with, respectively, the ‘old’ (pre-teleportation) and the ‘new’ (post-teleportation) maps. **C & D. Simulations of CANN model:** same analysis as in panels A & B, computed over 15 simulated teleportation events, with the same trajectory of the rodent as in the experimental data. Model parameters: *N* = 400 neurons, *γ*_*J*_ = 0.0025, *γ*_*V*_ = *γ*_*PI*_ = 0.4, *γ*_*W*_ = 6.25, *β* = 15, see Methods, Supplementary Text, and Supplementary Information Fig. 2&3 for detailed discussion of the choice of parameters.

In summary, the output of our map decoder, *m*_*t*_, can be interpreted as the correct cognitive state to read the positional code, see Fig. 2A. Even in the presence of fast dynamical flickers of the cognitive map, the location of the animal is robustly and coherently represented at all times. Our findings show that the hippocampus representation encodes both positional and contextual information in an independent and accurate way. Interestingly, the positional error computed with the map opposed to the decoded one (orange line in Fig. 3A) shows a significant reduction in the first seconds after the light switch; this non-trivial effect will be explained in detail in the next sections.

## Continuous attractor neural network model for the interplay between path integrator, visual cues, and memory

The findings above suggest that the stream of positional information to the hippocampus is maintained despite the presence of rapid changes of cognitive representations following the abrupt modification of visual cue (V) after the light switch. A natural hypothesis is that the path-integrator (PI) sends to the hippocampus information relative to the position in the ‘old’ map [11, 31], competing with the visual cue input associated to the ‘new’ map. To formalize this assumption, we introduce a continuous attractor neural network (CANN) model that contains the minimal ingredients to understand the effect of conflicting PI and V stimuli onto the hippocampal activity. In the CANN paradigm for memory storage and retrieval of cognitive maps [15–17, 36], the animal location at a certain time is represented as a self-sustained bump of neural activity. The bump is localized in the current position within a two-dimensional manifold, where place cells are embedded according to the positions of their place field centers in the real environment. We generalize this classical model by including two informational inputs on the memory network, from allothetic (visual cues) and idiothetic (path integrator) stimuli. The proposed interaction model is composed of the following four ingredients, see Fig. 4A and Methods:

a. A CANN *memory*, including excitatory recurrent connections of strength *γ*_*J*_ and global inhibition, designed to store and support two cognitive maps *m*, denominated *A* and *B*, which mimics the status of the CA3 place-cell network after learning of the two ‘environments’. This model of stochastic neurons was described and studied in [19, 36, 37].
b. A *visual-cue input* of amplitude *γ*_*V*_ onto place cells whose place fields match the current position of the rat, **r**_*t*_, in the cognitive map corresponding to the external light cue. The latter is denoted by *V* = *A* or *B*.
c. A *path-integrator input* of amplitude *γ*_*PI*_ that projects onto place cells whose place fields match the current position in the cognitive map corresponding to its own internal cognitive state [11], denoted by the variable *PI* = *A* or *B*.
d. An *effective feedback* from the hippocampal network to the path integrator, which stochastically maintains coherence between the recalled cognitive map and the path-integrator state.

**Fig. 4:**
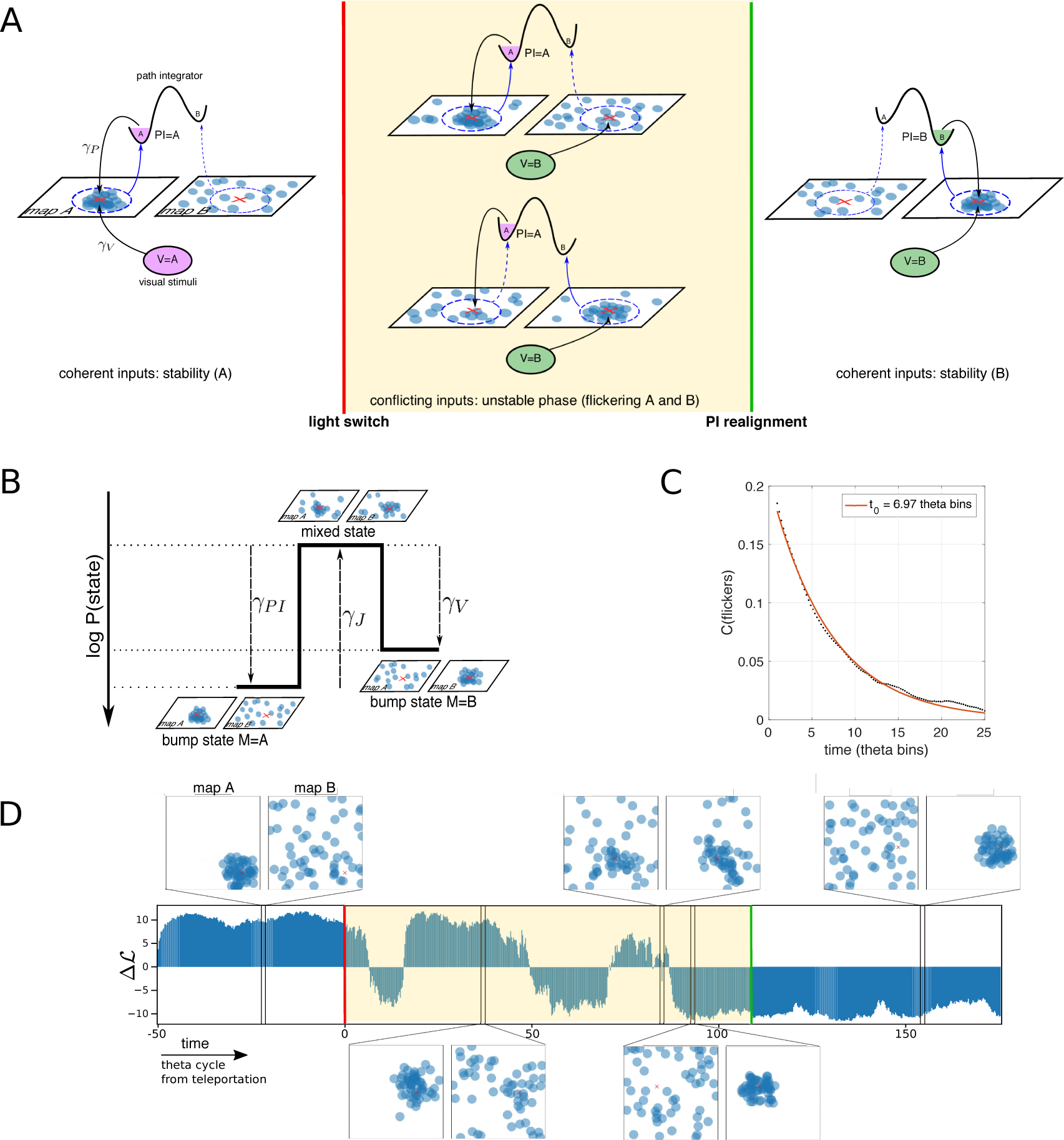
CANN model for interplay between path integrator, external stimuli, and memory. **A. Phenomenology of the CANN model**. The model is composed of a recurrent hippocampal network that has memorized two cognitive maps (place fields dispositions) denominated *A* and *B*, a path integrator input (PI), and a visual input (V). In the left panel both PI and V are activating place cells whose place-field centers (shown by blue dots) correspond to the position of the rodent (red cross X) in the cognitive map *A*. The activity is said to be localized in a bump around X in t map *A*, while it appears as sparse and uninformative if interpreted with respect to the place-field locations in map *B*. A feedback projection (blue arrows) from the hippocampal state (bump) to the path-integrator state (purple) maintains the stability of the system by enforcing that the retrieved hippocampal map and the PI state agree. After the light conditions have been switched (teleportation, red line), V projects on place cells encoding position X in map *B*. The two hippocampal cognitive maps are therefore in conflict, and the bump of activity is alternatively localized in *A* (center-top) or in **B** (center-bottom). When the hippocampal activity is localized in the cognitive map *B*, the feedback projection tries to realign the internal state of PI along the corresponding map. Once the reaglinment has succeeded (green line), both inputs are back to a coherent state, and stability is reached in the cognitive map relative to the post-teleportation external light conditions. **B.** Representation of the effective model for the activity bump and effects of parameters. The input strengths, *γ*_*PI*_ and *γ*_*V*_, contribute to push the hippocampal activity towards the corresponding cognitive states. Increasing the strength of recurrent connections, γ_*J*_, results in an effective barrier separating the two collective hippocampal states, giving rise to well formed bumps in either map *A* (left) or in map *B* (right). Due to this effective trap the bump state remains localized in either map for more than a single theta bin. **C.** Temporal correlation of flickering events (theta bins with cognitive state opposite to external light conditions) decay over c.a. 7 theta bins in simulated data. Same model parameters as in Fig. 3. **D.** Time trace of the log-likelihood difference *ΔL* (Eqn. (13) in Methods) in a simulated teleportation session; flickers can be observed during the conflicting phase following teleportation (shaded region). Prior to teleportation, and after the PI is realigned, inputs are coherent, and the system is stable: the sign of *ΔL* is constant, mirroring the localization of the bump in one map. Screenshots of the activity projected on the two cognitive maps are shown for five different times. From left to right: bump localized in map *A*, bump localized in map *A* during the conflicting phase, mixed state during the conflicting phase, bump localized in map *B* during the conflicting phase, bump localized in map *B*. The video of the simulation can be found in SI.

From a functional point of view, the CANN model mostly behaves, for a fixed position r of the rodent, as an effective two-state model for the hippocampal activity, as sketched in Fig. 4B. These two states correspond to the activity localized in map *A* or *B*; their probabilities are controlled by the intensity of, respectively, the path-integrator and visual-cue inputs. Note that the emergence of two well separated collective states from the microscopic CANN model is intrinsically due to the presence of recurrent connections, see Fig. 4B; A characterization of the effective barrier between the states is reported in SI, see Supplementary Fig. 1. The height of the barrier, controlled by the parameter *γ*_*J*_, and the amount of stochasticity in the individual neural dynamics are crucial ingredients to determine the dynamics of the model. In particular, these variables control the the time-correlation of flickers (Methods); For the chosen simulation parameters, the time correlation decays over ~ 7 theta bins (Fig. 4C) in accordance with data (Fig. 2C).

The typical outcome of a simulated experiment is shown in Fig. 4D. During the exploration phase preceding the light switch, the visual (b) and path-integrator (c) inputs jointly contribute to the stability of the internal representation of the position. A localized bump of activity, sustained by the recurrent connections (a), can be observed in the pre-switch map (Fig. 4A, left), say, *m* = *PI* = *V* = *A*. Right after the switch, the hippocampal network receives conflicting streams of information: *PI* = *A* differs from *V* = *B*. The path integrator is still activating place cells coding for the current position of the animal in the ‘old’ map, while the visual stream points to neurons coding for the same position in the ‘new’ map (Fig. 4A, center). This results in a *conflict* between the two bump representations, which are mutually incompatible due to the orthogonality of global remapping. Flickering is produced as an alternance between these two possible states, *m* = *PI* = *A* and *m* = *V* = *B*. During this conflicting phase (Fig. 4D, red region), the feedback (d) from the memory network to the path integrator tries to achieve coherence between the hippocampal and path-integrator states. When the bump is in the visually-driven, post-switch map (*m* = *B*), incoherence is strong, and the path integrator is more likely to be reset. Realigning the path integrator state with the external cue, *PI* = *V* = *m* = *B*, brings the conflict phase to an end, and the hippocampal state reaches stability (Fig. 4A, right).

Despite its conceptual simplicity, the model shows a rich phenomenology and reproduces in a strikingly-accurate manner the results of the analysis of the CA3 teleportation recordings. In Fig. 4D we show a representative time trace of the log-ratio *ΔL* (Eqn. (1)) in a simulated teleportation session. Alternate intervals of positive and negative *ΔL* signal the presence of map instability, as in [27], following the light switch (red vertical lines) and the path-integrator realignment (green vertical lines). Applying to the simulated data the same positional-error analysis as for the recorded data (Fig. 3A&B), we observe the same qualitative picture, see Fig. 3C&D. In particular, position is coherently encoded in the recalled cognitive map at all times.

## Flickering frequency is constant throughout the conflicting phase, whose duration is exponentially distributed

Our model predicts that (1) the duration of the conflict phase, i.e. the time elapsed from a light switch to the subsequent PI realignment, is exponentially distributed (Fig. 5A, top panel); (2) during the conflict phase, the flickering frequency i.e. the percentage of theta bins identified as flickers, is constant and independent of time (Fig. 5B, top panel).

**Fig. 5:**
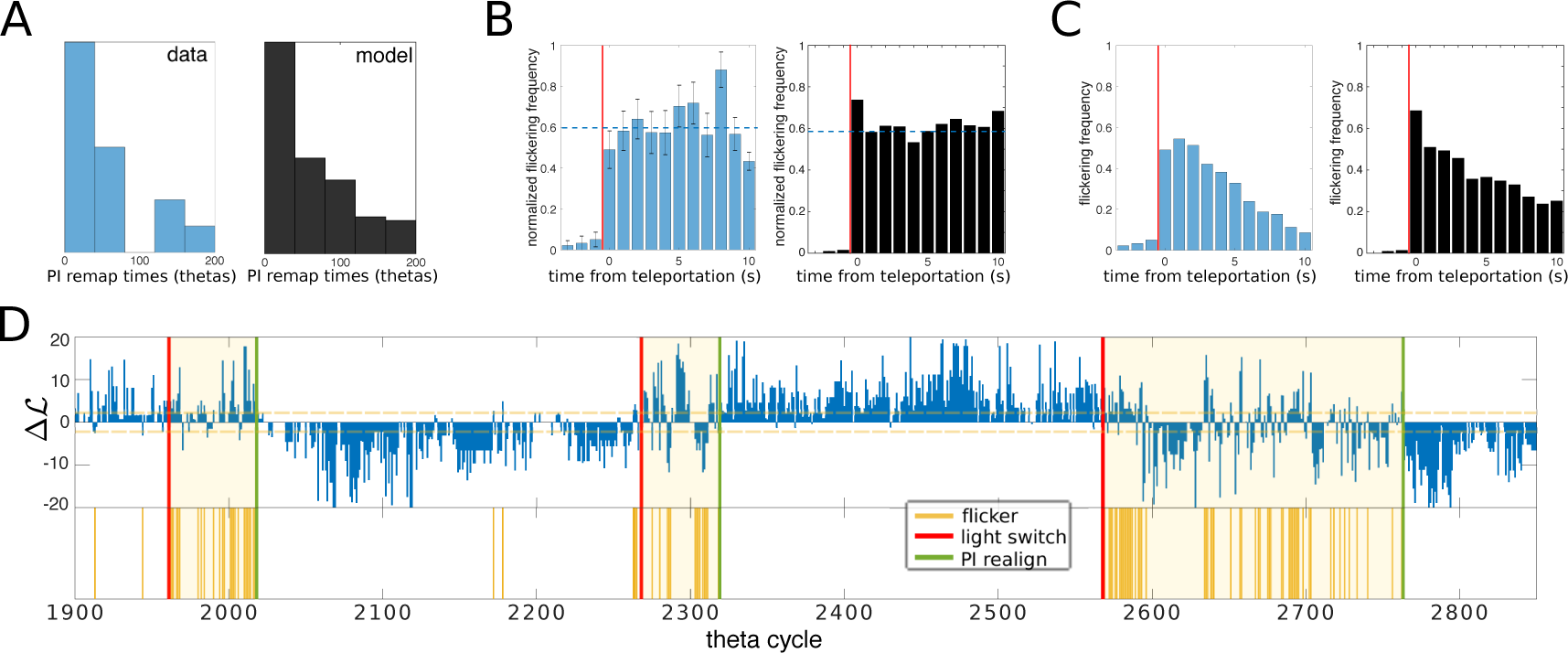
Flickering rate is homogenous within each conflicting period. **A.** Distribution of the path-integrator realignment times in a simulated session (top) and inferred from the recorded data (bottom). **B.** Mean Frequency of Flickers (MFF) during the conflicting phase, binned over 8 theta bins (~ 1 second) intervals. MFF is constant during the conflicting phase, both in the model (top) and in recordings (bottom). Realignment times inferred from data were obtained with a Bayesian procedure (Methods); histograms show flickering frequency in each time bin *t* normalized with respect to the fraction of conflicting phases, out of 15 teleportation events, that survived at least up to time *t*. **C.** Convolution of the two distributions shown in panels A and B, i.e. the MFF computed on the full test session, shows an apparent exponential decay both in the model (top) and in data (bottom, similar to the analysis shown in [27]). **D.** Map decoder output *ΔL* and inferred Pl-realignment times for the experimental test session; 3 out of 15 teleportations are shown. Light switches are marked with red lines, inferred Pl-realignment times are marked with green lines. Identified conflicting periods are shaded. Simulated data were obtained with the same model parameters as in Fig. 3.

In order to test these two predictions on CA3 recordings, we introduce a method to disentangle the flickering dynamics of the cognitive map and the realignment of the PI, the latter bringing an end to the former. We first infer the most likely PI-realignment time for each light-switch event, given the sequence of identified flickers (Methods). The outcomes are shown as green lines in Fig. 5D, and correctly separate conflicting phases (rich in flickering events) from coherent periods (during which the hippocampal representation is much more stable). The distribution of conflicting phase durations is approximately exponential in agreement with model prediction (1), with decay time *τ* = 53 theta bins (Fig. 5A, bottom panel). Dividing the test session into conflicting and coherent phases, we compute the frequency of flickers in the conflicting phase only. Consistently with the model prediction (2), the frequency of flickers is independent of the delay after the switch, with about 60% of theta bins in the conflicting phase carrying flickers (Fig. 5B).

Similar frequencies of flickers, close to one half, are obtained in the model when the two inputs have comparable strengths (*γ_PI_* ≃ *γ_V_* in Fig. 4B, see also Supplementary Fig. 3). A testable consequence of this balance is that the distributions of the sojourn times (durations of the periods in which the neural activity persists in a cognitive map, see Methods) in map A and in map B are similar. This prediction is confirmed by a further analysis of the CA3 recordings: the two distributions of the sojourn times are both exponential, with roughly the same decay times (Supplementary Fig. 5). This common time scale is related to the correlation time of the flickers (Fig. 2C & 4C), see Methods.

The combination of properties (1) and (2) explains the exponential decay in the frequency of flickers with the delay after the switch reported in [27] and [38]. While the frequency of flickers is constant and large in the conflicting phase, and constant and very low in the coherent phase, the duration of the conflicting phase is exponentially distributed. Hence the frequency of flickering theta bins, irrespectively of the phase, shows the same exponential decay, see Fig. 5C (top panel: simulated experiment, bottom panel: analysis of CA3 recordings). A detailed analysis of the data provides overwhelming statistical support to our two-fold explanation compared to a simple exponential decay of the flickering frequency (logarithmic likelihood-ratio test ~ 150, see Methods).

## Neural encoding of position reflects the presence of input mismatches

Our model allows us to better understand the subtle differences between the neural encodings of position in the conflicting and the coherent phases. In the latter phase, both path-integrator and visual inputs point to the neurons with place fields overlapping the rodent position **r** in map. During the conflicting phase, the two inputs excite the two place-cell populations centered in **r** in their respective maps, respectively, *m* = *PI* and *m* = *V*. Hence, while the bump of activity is mostly localized in one of the two maps (varying over time), some dispersion may be expected due to these incoherent inputs.

Mixed activity states, in which two (distinct) populations of neurons encoding the same position in the two maps are active, can be occasionally observed in the snapshots of the simulated activity in Fig. 4D, e.g. around theta bin *t* = 80. The overdispersion present during the conflicting phase has two consequences. First, the accuracy in position encoding is expected to be lower in the conflicting phase that in the coherent phase, see Fig. 6A, left panel. Secondly, the loss in accuracy is not due to some random noise in the neural activity, but to a transient bump-like activity in the ‘wrong’ map, opposite to the decoded one. This effect is clearly seen when we choose the opposite map to infer the rodent position. While this choice leads to very poor prediction during the coherent phase, the positional error is significantly reduced during the conflicting phase (Fig. 6B, left panel).

**Fig. 6:**
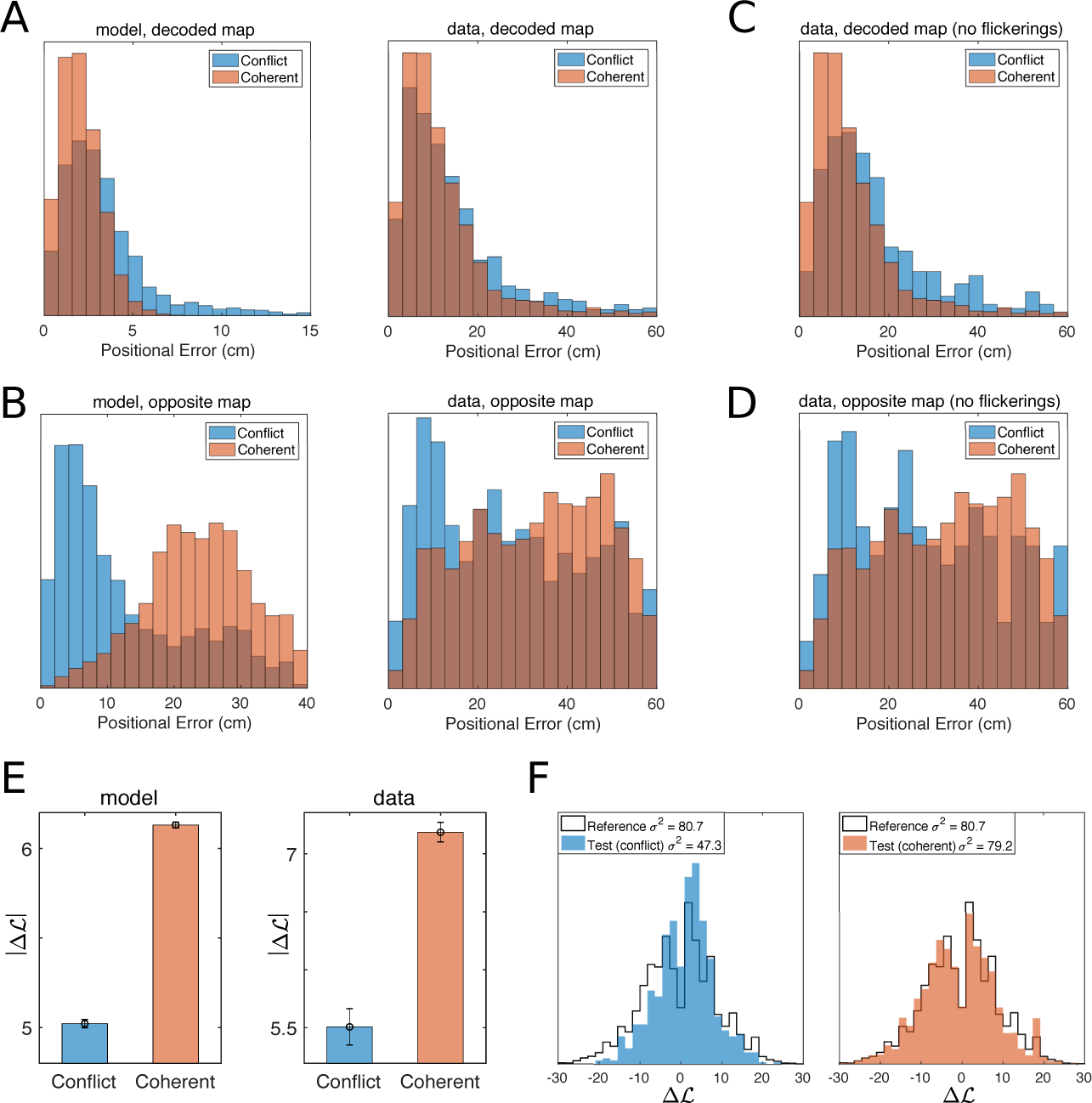
Distributions of positional errors in conflicting and coherent phases. **A.** Distributions of positional errors in the decoded cognitive map during coherent/stable (red) and conflicting/unstable (blue) phases. The model predicts (top panel) that the mean error is significantly increased during conflicting periods, mirroring the dispersion of the neural representation induced by the conflicting input streams. The same phenomenology is observed in recordings (bottom panel). **B.** The over-dispersion of the neural code in the ‘correct’ cognitive state (reported in panel A) is caused by an increased precision of the positional representation in the ‘opposite’ map, i.e. the one where the bump is not localized. This effect is significant in the model, top panel, and in the recorded data, bottom panel. **C, D.** Same analysis after exclusion of flickering theta bins. **E.** The absolute value of the log-ratio *ΔL*, Eqn. (1), is significantly reduced during the conflicting phase, both in the model (top) and in the data (bottom; ANOVA *p <* 10^−15^; conflicting: 5.51 ± 0.16 SEM, coherent: 7.19 ± 0.08 SEM). | *ΔL*| is a proxy for the completeness and stability of the bump (Methods). **F** Distributions of *ΔL* in the test session during the conflicting (left, blue) and the coherent (right, red) phases, compared to the distribution in the two reference sessions (black contour, same as in Fig. 2A).

To test these two predictions in CA3 recordings, we combine our positional analysis and our PI-realignment time inference procedure. In Fig. 6A (right), we compare the distributions of positional errors computed with the decoded map (acccording to the sign of *ΔL*) during conflicting and coherent phases (blue and red, respectively). Consistently with the model predictions, the positional error is significantly increased during the conflicting phase (ANOVA *p* < 8 × 10^−8^; conflicting: 14.7 ± 0.5 SEM, coherent: 12.3 ± 0.1 SEM). When computed with the opposite map, the positional error is obviously much higher than its counterpart computed with the decoded map, but a substantial decrease is found in the conflicting phase compared to the coherent phase, see Fig. 6B, right panel (ANOVA *p* < 5 × 10^−23^; conflicting: 27.8±0.6 SEM, coherent: 34.2±0.2 SEM), in full agreement with the model prediction. This effect also explains the relatively low value of the positional error obtained with the opposite map right after the switch, i.e. deep into the conflicting phase, compared to later times, see Fig. 3A&C. While this phenomenology is clear, it could in principle be affected by the presence of visual inputs projecting onto place cells during flickering events, i.e. when the ‘opposite’ map agrees with the external cues. In order to analyze the effect of the path integrator alone, we have restricted the analysis to theta bins whose decoded maps agreed with the visual inputs, i.e. to non-flickering theta bins. Results, shown in Fig. 6 C&D, are still statistically significant and in strong agreement with the model predictions (decoded map: ANOVA *p* < 1 × 10^−12^; conflicting: 17.1 ± 0.7 SEM, coherent: 12.3 ± 0.2 SEM; opposite map: ANOVA *p* < 1 × 10^−7^; conflicting: 28.9 ± 0.95 SEM, coherent: 34.2 ± 0.3 SEM). Our findings are robust against changes in the statistical threshold Lo for map decoding in the identification of conflict/coherent phases (Methods), see Supplementary Fig. 9.

The over-dispersion of the neural bump during the conflicting phase can also be observed from the reduction in (the absolute value of) the log-ratio, | *ΔL*|, see Eqn. (1). This quantity can be interpreted as a proxy for the completeness of the bump in one single map (Methods), larger | *ΔL*| corresponding to large bumps in either of the two maps and randomly scattered activity in the other map (Fig. 4A&D).

We find that the absolute value of *ΔL* is significantly reduced during the conflicting phase in CA3 data, see Fig. 6E (left panel, ANOVA *p* < 10^−15^; conflicting: 5.51 ± 0.16 SEM, coherent: 7.19 ± 0.08 SEM). Figure 6F shows the bimodal nature of the distributions of *ΔL* in the conflicting and coherent phases. While the reference and coherent-phase distributions coincide, the conflicting-phase distribution is more narrow, due to the overdispersion of the bump (reference *σ*^2^ = 80.7, coherent *σ*^2^ = 79.2, conflict *σ*^2^ = 47.3). This result provides further evidence for the predictive power of the CANN model.

## 3 Discussion

Our statistical inference-based data analysis allows us to quantify how well the CA3 neural activity encode various cognitve maps, and the position therein. Correlation-based procedures, e.g. used in [27], decode the cognitive state by comparing the instantaneous population activity to the average activity recorded in reference sessions at the same position of the rodent. Our functional-network based map decoder, instead, relies on the fact that the joint pairwise spiking activity of neurons is a fingerprint of the cognitive map [28, 34]. It does not need any knowledge of the sensory correlate (here, position), and could be used to decode generic brain states in other areas. The fast dynamics of cognitive maps studied in [27] and here results from an unrealistic sensory situation. Imposing artificial conflicts between inputs and studying their consequences is a standard approach to unveil the circuitry underlying the processing of multimodal sensory information in the hippocampus [25, 26] as well as in other brains areas, see for instance [39] for an illustration in the primary visual cortex where mismatches involve sensory and motor inputs. However, fast retrieval of functionally relevant maps, characterized by grouping and cognitive control, has also been observed in realistic settings, in which a behaving animal is required to maintain representations of two distinct spatial frames [20].

The position of the animal was accurately inferred at all times from the spiking activity using the place fields of the retrieved cognitive map (Fig. 1B). As a main finding, we show that the hippocampus maintains high-quality encoding of the position even if the contextual variable undergoes fast dynamical changes. This is explained in the model by the fact that inputs point to place cells coding for the physical position in both competing maps (Fig. 4A), and that the bump of activity is most often localized around these place cells in one map, and scattered all over the other map. Similar findings were reported in [27] (main text, Fig. 3d and Supplementary Information Fig. 8), within a statistical framework assuming a priori the consistency of positional representation during flickering events, as the cognitive map was decoded by comparing the neural activity to the mean-activity vectors at the recorded real position of the rat. The emergence of unambiguous, non-mixed representations was also underlined in [27], and shown to take place in the second half of the theta cycle. However, the detailed analysis of the CA3 recordings and of the model data shows a loss of quality of the bump state (reduction in absolute value of log-ratio |*ΔL*|) and an increased quality of position decoding in the opposite map (Fig. 6), providing evidence for the presence of partially mixed states.

Our model for the retrieval of hippocampal cognitive maps in the presence of inputs from the path integrator and visual cues is based on CANN theory [32, 40]. Two-dimensional CANN attractors, were previously applied to networks of place [15, 17] and grid [41, 42] cells. Indirect experimental evidence supporting CANN is now accumulating in various animals and brain areas. Evidence of a ring-shaped attractor region associated to head direction representation was recently reported in the drosophila central brain [43]. Attractor dynamics has also been associated to behavioral observation in a study on the monkey prefrontal cortex [44]. As for space representation, experimental support for attractor behaviour has been found in hippocampal CA1 [45] as well in grid cell [46] recordings. Further indirect evidence is provided by the pattern of connectivity in CA3, compatible with its functional role as an auto-associative attractor network [13], as suggested long ago based on anatomical and computational considerations [12, 47], and by the active nature of dendrites of mEC neurons, which enhances the robustness of attractors under environmental changes [48].

The detailed analysis of the CA3 recordings done here provides another indirect support for CANN theoretical framework, when multiple (two) cognitive maps are memorized. Memorization of the two attractors is obtained, in the model, by adding the corresponding connectivity matrices into the unique CANN connectivity matrix [16–19]. A detailed theoretical study of the mechanisms for transition from map to map was obtained in the absence of inputs, i.e. for spontaneous transitions induced by neural noise only [36]. A similar picture is found here in the presence of visual-cue inputs pointing to the ‘new’ map, while the path-integrator inputs point to the ‘old’ map in the conflicting phase. As inputs are of comparable magnitude, no single map is favored. The stochastic fluctuations resulting from the noise of the individual neurons are sufficient for the system to cross the activation barrier between the two memory states (maps) of the network, see Fig. 4B and Supplementary Fig. 1. The hippocampal network jumps intermittently from one cognitive map to the other, reproducing the flickering events experimentally identified and described in [27]. Transition rates between the two maps increase with the neural noise, modeled here by the parameter *β*, see Eqn. (12) in Methods and Supplementary Fig. 4. Neural noise relative to the population activity could also be effectively increased through the introduction of periodic (theta and gamma) modulations of the activity into the model [31, 37, 49]. The presence of rhythms is known to facilitate memory formation and integration of information [50, 51]. While theta oscillations can help produce flickering events as previously reported [31, 38], our work shows that such periodic modulations are not necessary. Transitions could also be facilitated by particular ‘confounding’ landmarks or positions in space, where the maps happen to be locally similar [16, 36].

The present model reproduces accurately all the observed flickering properties, without any need for a post-learning short-term plasticity of the CA3 network hypothetized in [38]. In particular, our model predicts that the flickering frequency is independent from the time spent after the teleportation event in the conflicting phase (Fig. 5B). This finding is at first sight in disagreement with the exponential decay of the flickering frequency reported in [27, 38]. However, the latter was obtained as a result of an averaging over many teleportation events. For a single event, accurate data analysis shows that our constant flickering rate hypothesis, when combined with the exponentially distributed realignment time of the path integrator (Fig. 5A), is much more likely than an exponential decreasing scenario.

Our model is based on the existence of two streams of inputs conveying, respectively, external landmark and self-navigation information. Recent studies have pointed to the grid cells network in mEC as the possible region that supports path integration, as their firing patterns are maintained in the dark [52], and the relative phases of grid cells seem to be largely unaffected by global remapping between environments of similar shapes [11, 53]. CANN-based approaches have been proposed to model grid-cell networks [41, 42], differing from hippocampal CANN mostly by the short-range nature of the inhibitory couplings. In much the same way the microscopic hippocampal CANN proposed here can effectively be reduced to a 2-state model (Fig. 4B), we expect CANN models for the grid-cell networks to be approximately described by a 2-state model, corresponding to the PI aligned with map A or B [11, 54, 55]. This motivates the simple model for the PI we have considered here.

In addition to sending projections towards the hippocampus, our model PI receives a feedback from the CANN, greatly increasing the probability of transition to the state agreeing with the instantaneous cognitive map [32, 56, 57]. Eventually, the state of the PI is realigned along the visual cue inputs, which stops the conflicting phase. Our model effectively implements a ratchet mechanism, locking the system into the coherent phase after a conflicting transient. Realignment of the path integrator based on visual landmarks is an important functional property, intended to limit the accumulation of errors in position estimation [58], and observed for large mismatch between external and internal inputs [24]. From a physiological point of view, projections exist from CA1 to mEC [32], and have been shown to be important for the formation of grid cells [57]. Hence, the feedback from the CANN, thought here to model CA3 activity, to the PI should be understood as effective.

As recently reported in [53], the impairement of the mEC grid firing resulted in a loss of path integrator in behaving rodents. As in our model, the recall of the pre-teleportation map, and, therefore, the whole flickering phenomenology are driven by the input stream from the path integrator to the CA3 network, we conjecture that flickering instabilities would disappear upon grid-cell impairement. Simultaneous recordings of mEC and CA3, as in [11], would be extremely useful to test our predictions and better describe the effect of the path integrator on the cognitive status of CA3.

## Acknowledgments

We are grateful to K. Jezek for providing us with the data of [27] and for fruitful discussions. We thank C. Schmidt-Hieber for insightful discussions and comments. We thank F. Rizzato, J. Tubiana, and S. Wolf for critical reading of the manuscript.

## Methods

### Cognitive map decoding from neural activity

Theta bins are identified with the Hilbert transform procedure of [27]. The activity of the *N* recorded neurons is binarized into each theta bin *t*: *s*_*i,t*_ = 1 if neuron *i* is active in bin *t*, 0 otherwise. For each cognitive map *m* = *A, B* a *Ising-model* probability distribution *P*_*m*_(**s**) for the activity configurations **s** = (*s*_1_, *s*_2_,…, *s*_*N*_) is inferred,

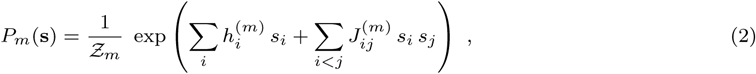

where *Z*_*m*_ is a normalization constant. Couplings (*J*^*m*^) and fields (*h*^*m*^) are determined such that the pairwise correlations and average activities in the neural population computed from *P*_*m*_ match their experimental counterparts in the reference session of environment *m*. These *inverse Ising problems* are solved using the Adaptive Cluster Expansion algorithm [59—62]. The inferred models (2) are then used to dynamically decode the map *m*_*t*_ during the test session (s) [28], based on the log-ratio of the probabilities of the activity configuration in time bin *t* in the two enviromments (main text Eqn. [1]), with the result

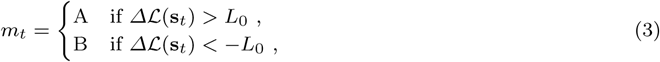

where the threshold *L*_0_ is chosen according to the required statistical confidence. We generally set *L*_0_ = log 10 ≃ 2.3.

After having decoded the map *m*_*t*_ in theta bin *t*, we define the flicker variable *f*_*t*_, equal to 1 if *m*_*t*_ does not match the light cue in theta bin *t*, to 0 otherwise.

### Temporal correlation of flickers and sojourn times

The time correlation of flickering events for delay *τ* is defined as

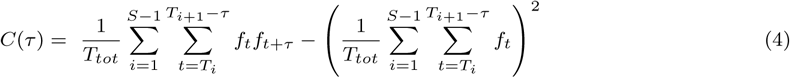

where *S* is the total number of switch events in the recorded data (*S* = 16 in [27]), *T*_*i*_ is the theta bin index of switch *i* (<*S*), and *T*_*tot*_ = *Ts* is the total number of theta bins in the test session. The time correlation *C*(*τ*) is typically exponentially decaying, with a decay time *τ*_0_, see Fig. 2C.

The correlation time *τ*_0_ is related to the sojourn time of the neural bump in the cognitive maps, defined as a sequence of contiguous theta bins decoded in the same map (Supplementary Information). Theta bins whose |*ΔL*| are lower than the threshold *L*_0_ are considered as belonging to the same map as the last statistically significant time bin. The distribution of sojourn times in each map is shown in Supplementary Fig. 5.

### Position decoding from neural activity

The arena is discretized into 60× 60 squared bins of 1 cm^2^ each, with integer coordinates (*x,y*) [27]. For each reference environment *m* ∈ (A, B) we construct the binary rate map, 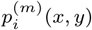, equal to the average of *s*_*i,t*_ over all theta bins *t* in which the rat is at position (*x,y*). Position is then decoded according to the naive Bayesian framework [63]: the probability of the activity configuration s_*t*_ *=* {*s*_1_, *s*_2_,…, *s*_n_} in theta bin *t* and at fixed position (*x,y*) reads

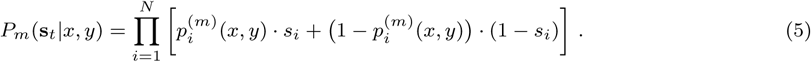

Once *m* is known, e.g. either through the map decoder or due to constant experimental conditions, the position of the rodent can be reconstructed from the recorded neural activity through

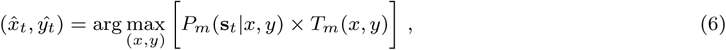

where the maximum is computed over the 60 × 60 possible positions. *T*_*m*_(*x,y*) is the number of theta bins spent by the rodent at position (*x, y*) during the reference session of map *m*; we use it as a prior to favor positions where the rodent is more likely to be, irrespectively of the neural activity.

### Continuous Attractor Neural Network model for hippocampal activity

The hippocampal population includes *N* place cells. For each cell *i* = *1…N* the place-field centers coordinates, 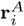 and 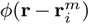, are drawn uniformly and independently at random in the squared environments, respectively, A and B. The linear size of each square is denoted by *L*.

Neural activities are represented by binary variables: *s*_*i,t*_ = 0 or 1 if neuron *i* is, respectively, silent or active in time bin *t* = 1, 2, 3,…. The duration of a time bin is the theta cycle over *K*; results reported here were obtained with *K* = 4, which corresponds to approximately 30 ms.

The total input received by neuron *i* at time *t* is

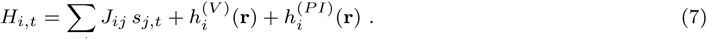

The three terms on the right hand side of Eqn. [7] represent, in order:

– the input due to recurrent connections in the hippocampal network. The underlying assumption is that connections have emerged from learning during the exploration of the two environments by the rodent: Place cells that turned out to be simultaneously active in either environment have developed positive couplings. Couplings are defined through

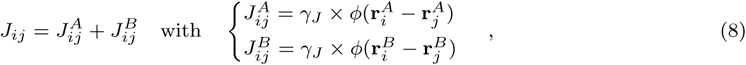

where *γ_J_* controls the strength of the connections, and

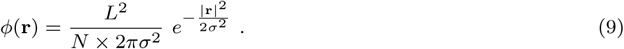

Parameter *σ* in Eqn. [9] is the spatial scale corresponding to the width of the place fields. The prefactor in Eqn. [9] ensures that *φ* is dimensionless and that, on average, the sum of the Gaussian factors 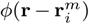 over all neurons *i* is close to unity for every possible position *r* and map *m*. the visual input

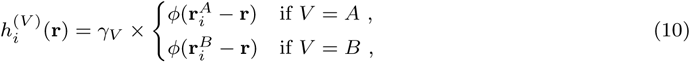

depends on the position **r** of the rodent. We again assume that, during the exploration of the environment, visual-cue projections onto place cells have been strenghtened through learning. For simplicity we use the same function *φ* as in the recurrent connections, see Eqn. (8), to characterize the portion of environment in which visual cues project onto a specific place cell *i*.

– the path-integrator input

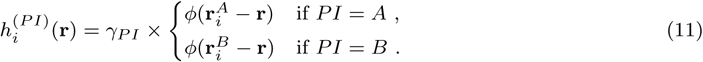

PI inputs have the same functional dependence over space as visual-cue related inputs. The amplitudes of both input types are tuned by the parameters *γ_PI_* and *γ_V_*.

All neurons undergo stochastic updating of their activities from time bin *t* → *t* + 1 according to their total inputs. The activity of neuron *i* at time *t* +1 is chosen to be

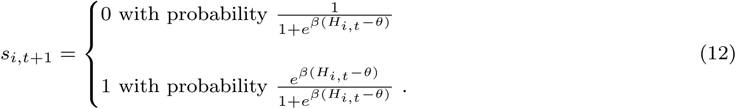

To enforce global inhibition in the population activity, the value of the threshold *θ* is dynamically adjusted so that an average fraction *f* of the neurons is active at any time. Parameter *β* controls the amount of noise in the neural dynamics. For *β* → 0 neuron activities are random and independent of their inputs, while, for *β* →∞, they deterministically follow the signs of the inputs (after substraction of the threshold *θ*). The average activity of cell *i* at time *t* + 1 is therefore a monotonously increasing sigmoidal function of its total input *H*_*i,t*_ at time *t*, with maximal slope equal to *β*/4 in *H*_*i,t*_ = *θ.*

The properties of this CANN model in the absence of any visual and PI inputs, i.e. for *γ_V_* = *γ*_*PI*_ = 0, were analytically studied in [19, 36, 37], see Supplementary Information for further discussion. The log-ratio *ΔL* defined in Eqn. (1) for the decoding of cognitive maps has a direct counterpart in our CANN model as the difference between the contributions to the log-probability of an activity configuration s when the bump is localized in maps A and B,

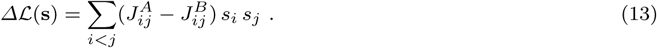

### Mechanism for path-integrator realignment

The path integrator is described as a two-state model, *PI* = *A* or *B*. Its dynamics is stochastic and Markovian: in each time bin *t*, the state *PI* can jump into state *PI*’ with transition probabilities *R*(*PI* → *PI*’), independently of the previous states. The feedback from the hippocampal network to the path integrator is expressed in the dependence of *R* on the hippocampal map *M*_*t*_ at time *t*. To favor transitions to the state *PI*’ agreeing with the current map *M*_*t*_, we introduce the following witness function for the presence of the bump in map *m* = A, B:

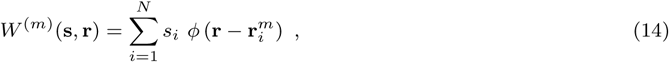

where **s** and **r** are, respectively, the activity configuration and the position of the rodent at time *t*. Due to the normalization of *φ* in Eqn. [9], we expect *W*^(*m*)^ to be close to one for the retrieved map *m* = *M*_*t*_ and to be much smaller for the opposite map.

We impose the preference for realigning the path-integrator state in accordance with the hippocampal map through the ratio between the two reciprocal transition probabilities,

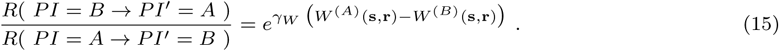

Here, *γ*_*W*_ is a positive parameter allowing us to tune the strength of the preference. If the hippocampal bump of activity is localized in, say, map *A*, the right hand side of Eqn. [15] will be strongly positive, and the probability of realigning the path integrator to *PI*’ = *A* will be much larger than the probability of the reciprocal transition. A solution to the constraint expressed by Eqn. [15] is given by

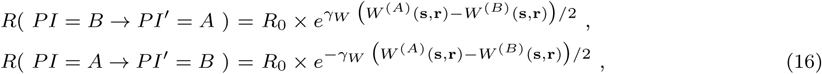

where *R*_0_ is a positive number. In the absence of bias (*γ*_*W*_= 0), the inverse of *R*_0_ may be interpreted as the average time scale between two realignments of the path-integrator state.

The model for the path-integrator dynamics is entirely defined by the transition probabilities in Eqn. [17] and the probability conservation identities:

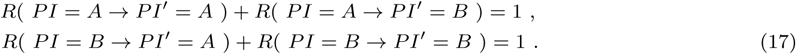

### Inference of path-integrator realignment times

Defining *τ* as the PI-realignment time, *t* = 0 as the time bin corresponding to the light switch and T as the time bin corresponding to the next switch (end of analyzed data), we assume the probability *p*(*t*) for time bin *t* to be a flickering event to be

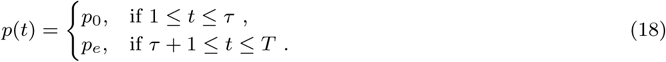

Here, *p*_0_ is the constant flickering probability, and *p*_*e*_ is the baseline decoding error, see Supplementary Information for discussion of the values of parameters *p*_0_ and *p*_*e*_.

We write the log-likelihood of the parameter τ as a function of the identified flickering sequence **f** = {*ft*} as follows:

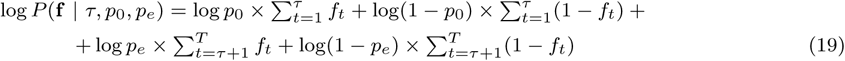

We then maximize this log-likelihood over *τ* to infer the most likely value *τ*\* of the realignment time. The procedure is repeated for all light switches.

### Independence of frequency of flickers from delay after light switch

We consider two hypothesis:

a. *H*_*decay*_ = the flickering probability depends on time as a decaying function that can be inferred from data, that can be inferred from the full test session (15 lght switches)
b. *H*_*constant*_ = the flickering probability is constant throughout the conflicting period of varying duration, which can be inferred from data (see previous section).

Following the Bayesian information criterion [64], we parametrize each model with the same number of variables. For hypothesis *H*_*decay*_, we estimate the flickering frequency as a function of time from the average frequency computed over the full test session (Fig. 5C, bottom), in bins of one second-width (8 theta cycles), up to 15 seconds after the light switch. For later delays (>15 s) the flickering frequency is set to a baseline error probability, *p*_*e*_ = 0.01. For hypothesis *H*_*constant*_, we infer the most likely PI realignment times for each one of the 15 light-switch events (see Section above). The associated flickering probability *p*_*t*_ is then set to *p*_0_ = 0.55 until the inferred PI realignment time *τ*\*, and equal to the baseline probability *p*_*e*_ = 0.01 afterwards. We then compute the likelihoods of both hypothesis given the observed data (identified flickering theta bins **f**) through

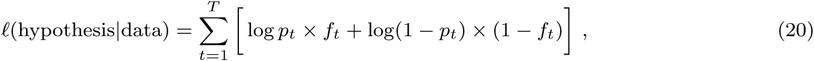

where *T* is the total length of the test session. The above expression is then summed over all light-switch events. We define the difference of the two log-likelihoods as

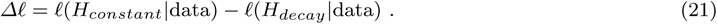

The constant flickering frequency hypothesis *H*_*constant*_ is extremely more likely (Δ*l* ~ 150) than the decaying model *H*_*decay*_. The result is robust against changes in the parameters, see Supplementary Fig. 6 and Supplementary Information (Detailed Methods).

